# A dynamic template complex mediates Munc18-chaperoned SNARE assembly

**DOI:** 10.1101/2022.09.09.507208

**Authors:** Jie Yang, Huaizhou Jin, Yihao Liu, Yaya Guo, Yongli Zhang

## Abstract

Munc18 chaperones assembly of three membrane-anchored soluble N-ethylmaleimide-sensitive factor attachment protein receptors (SNAREs) into a four-helix bundle to mediate membrane fusion between vesicles and plasma membranes, leading to neurotransmitter or insulin release, GLUT4 translocation, or other exocytotic processes. Yet, the molecular mechanism underlying chaperoned SNARE assembly is not well understood. Recent evidence suggests that Munc18-1 and Munc18-3 simultaneously bind their cognate SNAREs to form ternary template complexes - Munc18-1:Syntaxin-1:VAMP2 for synaptic vesicle fusion and Munc18-3:Syntaxin-4:VAMP2 for GLUT4 translocation and insulin release, which facilitate binding of SNAP-25 or SNAP-23 to conclude SNARE assembly. Here, we further investigate the structure, dynamics, and function of the template complexes using optical tweezers. Our results suggest that the synaptic template complex transitions to an activated state with a rate of ∼0.05 s^−1^ and ∼6.8 k_B_T higher energy for efficient SNAP-25 binding. The transition depends upon the linker region of syntaxin-1 upstream of its helical bundle-forming SNARE motif. In addition, the template complex is stabilized by a poorly characterized disordered loop region in Munc18-1. While the synaptic template complex efficiently binds both SNAP-25 and SNAP-23, the GLUT4 template complex strongly favors SNAP-23 over SNAP-25, despite similar stabilities of their assembled SNARE bundles. Together, our data demonstrate that a highly dynamic template complex mediates efficient and specific SNARE assembly.

**Significance:** Munc18-1 chaperones coupled folding and assembly of three synaptic SNAREs, syntaxin-1, VAMP2, and SNAP-25, into a four-helix bundle to mediate membrane fusion and neurotransmitter release. Recent evidence suggests that Munc18-1, syntaxin-1, and VAMP2 first form a weak template complex and then bind to SNAP-25 to complete SNARE assembly. However, the dynamics and function of the template complex are not well understood. Using optical tweezers, we found that the template complex undergoes a conformational change to bind SNAP-25 in a way dependent upon the syntaxin linker region and that Munc18 kinetically proofreads SNARE pairing not governed by its thermodynamic stability. Our study reveals a more dynamic template complex than that seen in its cryo-EM structure.

---

The fusion of synaptic vesicles with the pre-synaptic membrane leads to neurotransmitter release into the synaptic junction required for neurotransmission. The fusion process is mediated by VAMP2 (or synaptobrevin) anchored on the vesicular membrane (v-SNARE) and syntaxin-1 and SNAP-25 located on the plasma membrane (t-SNAREs) (Sollner et al., 1993). The three SNAREs fold and assemble into a parallel four-helix bundle in a coupled manner, drawing the two associated membranes into proximity to induce fusion (Gao et al., 2012; Sutton et al., 1998). The core of the four-helix bundle contains 15 layers of hydrophobic residues and a middle ionic layer with three glutamines and one arginine, designated as -7 - +8 layers from the N-terminus to the C-terminus, with the middle ionic layer as “0” layer. SNARE assembly is chaperoned by Munc18-1 and other regulatory proteins to achieve the speed and accuracy required for neurotransmission (Baker and Hughson, 2016; Jiao et al., 2018; Rizo, 2022; Shen et al., 2007; Shu et al., 2020; Sudhof and Rothman, 2009). Munc18-1 homologs (collectively Sec1/Munc18- (SM-) family proteins) are essential for all SNARE-mediated membrane fusion (Verhage et al., 2000; Zhang and Hughson, 2021). In particular, Munc18-3 and its cognate SNAREs syntaxin-4, VAMP2, and SNAP-23 mediate insulin release in β cells and fusion of glucose transporter 4 (GLUT4) containing vesicles with the plasma membrane for glucose uptake in muscle or fat cells (Thurmond and Gaisano, 2020; Yu et al., 2013). Malfunctions of the SNARE-SM fusion machinery have been linked to many diseases (Rebane et al., 2018; Stamberger et al., 2016; Thurmond and Gaisano, 2020). However, the molecular mechanism underlying physiological, Munc18-chaperoned SNARE assembly is generally not well understood.

Munc18-1 guides synaptic SNARE assembly via a series of intermediates different from those populated during spontaneous SNARE assembly (Ma et al., 2013; Zhang and Hughson, 2021). SNAREs alone assemble via the labile 1:1 syntaxin-1:SNAP-25 t-SNARE complex (Pobbati et al., 2006; Zhang et al., 2016a). Overall, the assembly occurs with a low speed and accuracy due to frequent misassembly into various fusion-incompetent byproducts (Brunger, 2005; Choi et al., 2018; Weninger et al., 2008). Furthermore, the t-SNARE complex is disassembled by the ubiquitous AAA+ ATPase NSF and its adaptor SNAP (Choi et al., 2018; Ma et al., 2013; Stepien et al., 2019). Thus, spontaneous assembly via a t-SNARE assembly intermediate is unlikely to occur *in vivo*. Over the past decade, an alternative Munc18-1-chaperoned SNARE assembly pathway has gained increasing experimental support (Jiao et al., 2018; Ma et al., 2013; Zhang and Hughson, 2021; Zhao et al., 2013). Munc18-1 tightly binds to syntaxin-1 to chaperone syntaxin-1 trafficking from the endoplasmic reticulum (ER) to the plasma membrane (Zhou et al., 2013). Munc18-1 binding also sequesters syntaxin-1 from associating with other SNAREs, forming closed syntaxin (Burkhardt et al., 2008; Misura et al., 2000). Munc13-1 or syntaxin-1 mutations help open the closed syntaxin to allow VAMP2 to bind, forming a tetrameric or ternary template complex in the presence or absence of Munc13-1, respectively (Shu et al., 2020). Using optical tweezers, we recently identified the synaptic template complex and characterized its stability and salient structural features (Jiao et al., 2018). We found that SNAP-25 quickly binds the template complex to conclude SNARE assembly and displace Munc18-1 from the four-helix bundle. This Munc18-1-dependent pathway is rapid compared with the spontaneous SNARE assembly under the same experimental conditions. Mutations that destabilize the template complex also impair SNARE assembly, membrane fusion, and/or neurotransmission. Finally, similar template complexes are found for SNARE-SM machines responsible for other membrane trafficking pathways, including the GLUT4 template complex Munc18-3:Syntaxin-4:VAMP2 (Baker et al., 2015; Jiao et al., 2018). Thus, the template complex is probably a conserved physiologically relevant intermediate that significantly enhances the speed and accuracy of SNARE assembly (Zhang and Hughson, 2021).

In a major step toward understanding Munc18-1-chaperoned SNARE assembly, Stepien et al. recently reported 3.5-3.7 Å-resolution cryo-EM structures of the synaptic template complex (Figure 1A) (Stepien et al., 2022). The structure confirms many features of the structural model derived from earlier single-molecule and structural studies: the N-terminal SNARE motifs of syntaxin-1 and VAMP2 are aligned as parallel helices, while the C-terminal halves are kept separated also in helical conformations (Baker et al., 2015; Jiao et al., 2018). In addition, the syntaxin region N-terminal to the SNARE motif, including the N-terminal peptide and the Habc three-helix bundle, stabilizes the template complex (Jiao et al., 2018). Importantly, the cryo-EM structure reveals a small four-helix bundle formed by the N-terminal regions of the syntaxin-1 and VAMP2 SNARE motifs and the syntaxin linker region (Stepien et al., 2022). The syntaxin linker region connects the Habc domain to the SNARE motif and, in the cryo-EM structure, folds into two α-helices (Fig. 1A, Hd and He helices). Surprisingly, this four-helix bundle and the entire N-terminal region of the VAMP2 SNARE motif minimally interact with the underlying Munc18-1 surface - the tip of the helical hairpin in domain 3a (or 3a hairpin). Consequently, the templated syntaxin-1 and VAMP2 associate with Munc18-1 mostly via the splayed C-terminal regions of their SNARE motifs. This structural feature contrasts with the previous observations that the tip of the 3a hairpin binds to SNAREs and is essential for neurotransmission (Baker et al., 2015; Munch et al., 2016). Therefore, the cryo-EM structure opens a new avenue to further test the dynamics and function of the template complex.

**Fig. 1.**
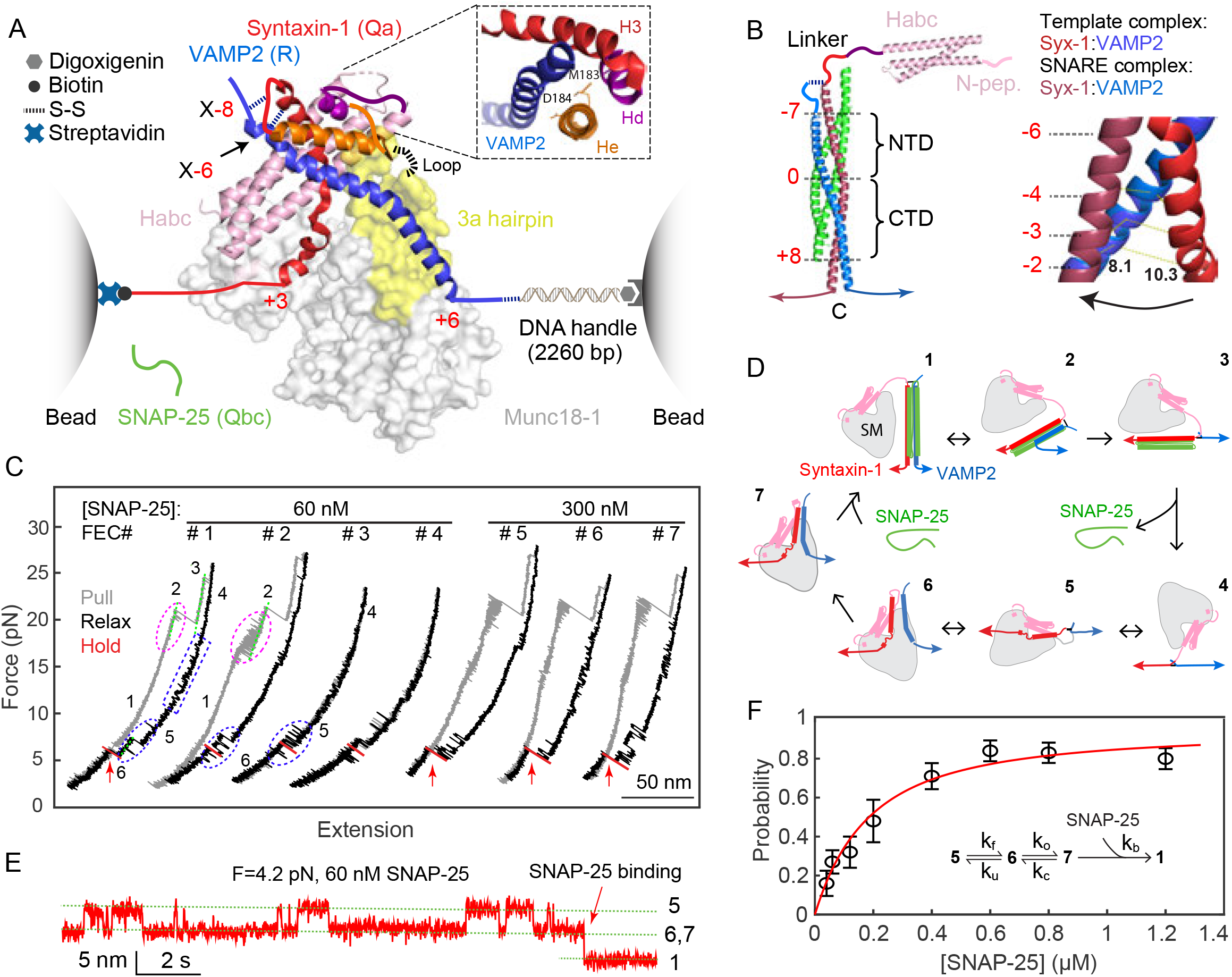
Single-molecule manipulation based on optical tweezers revealed an activated template complex nucleating SNARE assembly. (A) Experimental setup and cryo-EM structure of the template complex (PDB ID: 7UDB). Synatxin-1 and VAMP2 are crosslinked at either the -8 layer (X-8) or the -6 layer (X-6) as indicated. The 3a helical hairpin of Munc18-1 (yellow) is the primary VAMP2 binding interface, with the disordered loop region at the tip indicated by a dashed line. The inset shows the N-terminal four-helix bundle, with syntaxin-1 M183 and D184 in the He helix located in the binding interface. (B) Structure of the fully assembled SNARE complex containing the SNARE four-helix bundle (PDB ID: 1SFC) and the syntaxin Habc domain (PDB ID: 1BR0) (left panel) and different helical conformations of N-terminal SNARE motifs of syntaxin-1 and VAMP2 in the template complex and the SNARE four-helix bundle (right panel). The characteristic layers are labeled in red numbers. The middle ionic layer, or “0” layer, divides the SNARE four-helix bundle into the N-terminal domain (NTD) and the C-terminal domain (CTD). In the right panel, the distances between the two residues at the -2 layer are labeled. (C) Force- extension curves (FECs) obtained by repeatedly pulling (grey) and relaxing (black) a single Syx- VAMP conjugate or holding at a constant trap separation (red) in the presence of Munc18-1 and SNAP-25 in the solution. The states associated with different FEC regions are labeled (see D) and the SNAP-25 binding events are indicated by red arrows. (D) Diagrams showing different SNARE folding/assembly and Munc18-1 binding states and their transitions (black arrows): 1, the fully assembled SNARE four-helix bundle; 2, the half-zippered SNARE bundle; 3, the unzipped t- and v-SNAREs; 4, the fully unfolded SNAREs; 5, open syntaxin; 6, the inactive template complex; 7, the activated template complex. (E) Extension-time trajectories at constant trap separation or mean force showing reversible folding and unfolding transition of the template complex and irreversible SNAP-25 binding. (F) The probability of SNAP-25 binding to the template complex as a function of SNAP-25 concentration (symbol) and its best model fit (red curve). Each probability was derived from more than 30 independent measurements (N), with the arrow bars indicating the standard deviation (SD). The inset shows the reaction scheme and the associated rate constants (SI Appendix).

The formation of the template complex greatly enhances the specificity of SNARE pairing (Jiao et al., 2018; Koike and Jahn, 2022; Shen et al., 2007; Yu et al., 2013), which is otherwise promiscuous (Brunger, 2005; Yang et al., 1999). SNAREs are divided into Qa, Qb, Qc, and R-SNAREs based on the characteristic glutamine (Q) or arginine (R) residues in the ionic layer (Fasshauer et al., 1998). The template complexes identified so far are formed by SM proteins and their cognate Qa and R-SNAREs, or the corresponding syntaxin- and VAMP-family SNAREs (Fig. 1A) (Baker et al., 2015; Jiao et al., 2018; Zhang and Hughson, 2021). The formation of template complexes promotes specific pairing between Qa- and R-SNAREs. However, it is not known whether and how template complexes proofread the association of SNAP-25-family (Qbc) SNAREs, including SNAP-23, SNAP-29, and SNAP-47 in vertebrates (Kadkova et al., 2019), into SNARE four-helix bundles, as SM proteins do not seem to directly bind these Qbc-SNAREs. This question is physiologically and pathologically relevant because cognate SNARE-SM fusion machines responsible for many vesicle trafficking pathways are unclear. Identification of these machines is partly complicated by the fact that multiple SM proteins are often involved in the same trafficking pathways with distinct and overlapping sets of SNAREs. For example, Munc18-1, Munc18-3, Syntaxin-1, Syntaxin-4, SNAP-25, SNAP-23, and VAMP2 all participate in insulin release in β cells, leading to complex kinetics of insulin release and disease phenotypes (Thurmond and Gaisano, 2020). It has been proposed that these SM and SNARE proteins might cooperate in membrane fusion in a promiscuous manner. However, detailed characterizations of their interactions have generally been lacking.

It is technically challenging to study SM-chaperoned SNARE assembly (Zhang, 2017). The template complexes are marginally stable and have only been prepared with crosslinked SNAREs (Jiao et al., 2018; Stepien et al., 2022). It remains unclear how crosslinking affects the structure and dynamics of the template complex. Furthermore, the template complex is highly dynamic and can unfold into closed, open, or unfolded syntaxin conformations, partly due to multi-valent interactions between Munc18-1 and SNAREs (Zhang and Hughson, 2021). SNARE or Munc18-1 mutations may differentially affect these states, complicating data interpretation. Here, we used high-resolution optical tweezers to further dissect the stability, dynamics, and function of the template complex. We found that the template complex undergoes a conformational change to bind SNAP-25, which requires folding of the syntaxin linker region and SNARE binding of a poorly characterized Munc18-1 loop region. In addition, crosslinking affects the stability and SNAP-25 binding of the template complex. Finally, we showed that template complexes proofread the binding of Qbc-SNAREs.

## Results

### Template complex undergoes a conformational change to bind SNAP-25

The α-helices formed by the N-terminal region of the syntaxin SNARE motif in the template complex and the assembled four-helix bundle show different orientations relative to the corresponding VAMP2 helices (Fig. 1B). In addition, the syntaxin-1 and VAMP2 helices in the template complex gradually separate from the -6 layer to the -1 layer. Thus, the template complex must undergo a conformational change for SNAP-25 to bind and form the SNARE four-helix bundle. We hypothesized that this conformational change may become rate-limiting at a high SNAP-25 concentration. To test this hypothesis and detect the conformational change, we adopted our single-molecule optical tweezers assay for the Munc18-1-chaperoned SNARE assembly (Jiao et al., 2018; Shu et al., 2020). We crosslinked syntaxin-1 and VAMP2 at the -8 or -6 layer to form a Syx- VAMP conjugate, designated as X-8 or X-6, respectively, through a disulfide bridge. These crosslinking sites were strategically chosen to also open the closed syntaxin for template complex formation. The open syntaxin-1 presumably binds to Munc18-1 in a way to sequester its residues between +1 and +3 layers but expose most of its N-terminal SNARE motif, consistent with earlier structural and single-molecule studies (Eisemann et al., 2020; Jiao et al., 2018; Zhang and Hughson, 2021). The C-termini of syntaxin-1 and VAMP2 were connected to polystyrene beads either directly or via a DNA handle (Cecconi et al., 2005). The beads were held in two optical traps as force and displacement sensors. The extension and tension of the protein-DNA tether were measured to report the conformational changes of the SNAREs and their associated energies in the presence of Munc18-1 and SNAP-25 in the solution.

We first tested the construct crosslinked at X-8. When pulled to a high force, a single pre-assembled SNARE complex unfolded stepwise with characteristic intermediate states and kinetics as seen in the force-extension curve (FEC, Fig. 1C, FEC#1, grey curve) (Gao et al., 2012). These intermediates include a half-zippered SNARE bundle and the unzipped t- and v-SNAREs (Fig. 1D, states 2 and 3). During relaxation, Munc18-1 first binds to syntaxin-1, forming the open syntaxin in a reversible manner in the force range of 10-17 pN (Fig. 1C, blue dashed rectangle; Fig. 1D, state 5) (Jiao et al., 2018). At a lower force range of 3-7 pN, VAMP2 reversibly associates with open syntaxin to form the template complex (Fig. 1C, blue dashed oval; Fig. 1D, state 6). We then held the template complex around its equilibrium force with half unfolding probability and waited for SNAP-25 binding (Fig. 1C, red region, and Fig. 1E). The binding was identified as a sudden extension drop (Figs. 1C &1E, red arrows) and the resultant fully assembled SNARE complex was confirmed by subsequent pulling (Fig. 1C, FEC#2, grey curve). If no SNAP-25 binding was detected within 120 seconds, the Syx-VAMP conjugate was first relaxed to lower force and then pulled again. Finally, we repeated the cycle of relaxation to assemble the SNARE complexes and subsequent pulling to unfold the assembled SNARE complexes for tens of rounds in total on the same and different conjugates at each SNAP-25 concentration (Table 1). The probability of SNARE assembly per round as a function of SNAP-25 concentration was measured (Fig. 1F).

**Table 1.** Properties of the neuronal template complex crosslinked at X-8 or X-6 containing syntaxin-1 (Syx) or Munc18-1 (M18) modifications. The number in parenthesis is the standard error of the mean (SEM). N.D., not detected, which means that the stability of the template complex is too weak to be detected by our method, estimated to be <1.5 k_B_T. N.A., not available.

| Crosslinking site | Syx or M18 modification | Template complex |  | Open syntaxin<sup>b</sup> |  | SNAP-25 binding (60 nM) |  |
| --- | --- | --- | --- | --- | --- | --- | --- |
|  |  | Unfolding energy (k<sub>B</sub>T) | Equilibrium force<sup>a</sup> (pN) | Unfolding energy (k<sub>B</sub>T) | Equilibrium force (pN) | Prob.<sup>c</sup> | N<sup>d</sup> |
| **X-8** | WT | 5.2 (0.1) | 5.1 (0.1) | 2.6 (0.2) | 10.8 | 0.27 | 55 |
|  | Syx Δ147-197 | 4.7 (0.2) | 5.4 (0.2) | 2.7 (0.3) | 11.1 (0.4) | 0 | 18 |
|  | Syx Δ147-170 | 4.8 (0.2) | 5.2 (0.1) | 3.0 (0.2) | 10.3 (0.1) | 0.24 | 29 |
|  | M18 GS314-326 | N.D. | N.A. | 2.7 (0.2) | 10.6 (0.1) | 0 | 14 |
|  | M18 Δ314-326 | N.D. | N.A. | N.D. | N.A. | 0 | 22 |
| **X-6** | WT | 3.9 (0.1) | 5.8 (0.1) | 2.4 (0.2) | 11.4 (0.2) | 0.035 | 50 |
|  | Syx Δ147-197 | 3.5 (0.1) | 6.1 (0.1) | 2.4 (0.2) | 12.6 (0.2) | 0.029 | 11 |
|  | Syx M183A | 3.7 (0.2) | 6.2 (0.1) | 2.2 (0.2) | 11.5 (0.2) | 0.047 | 21 |
|  | Syx D184P | 3.5 (0.2) | 6.3 (0.2) | 1.9 (0.2) | 10.8 (0.3) | 0.056 | 18 |
|  | M18 GS314-326 | N.D. | N.A. | 2.3 (0.2) | 10.3 (0.3) | 0 | 19 |
|  | M18 Δ314-326 | N.D. | N.A. | N.D. | N.A. | 0 | 29 |

<sup>a</sup> Mean of two average forces for the unfolded and folded states when the two states are equally populated (Rebane et al., 2016). The equilibrium force of the template complex generally correlates with its unfolding energy.
<sup>b</sup> Detected as the syntaxin- and Munc18-1-dependent transition in the force range of 10-15 pN.
<sup>c</sup> Probability of SNAP-25B binding to the template complex per relaxation.
<sup>d</sup> Total number of relaxations to detect SNAP-25 binding.

Munc18-1-chaperoned SNARE assembly could be detected in the presence of 0.04 μM SNAP-25 with a probability of 0.16. Under this condition, spontaneous SNARE assembly in the absence of Munc18-1 barely occurred, confirming that Munc18-1 catalyzes SNARE assembly (Jiao et al., 2018). As the SNAP-25 concentration increased to 0.12 μM, the probability increased approximately linearly, consistent with rate-limiting bimolecular binding between SNAP-25 and the template complex. Interestingly, further increasing the SNAP-25 concentration led to a probability plateau at ∼0.8, suggesting that, at high SNAP-25 concentrations, a SNAP-25- independent step precedes SNAP-25 binding. We propose that, at all SNAP-25 concentrations, SNAP-25 binds solely to an activated template complex state that interconverts with an inactive state. At high SNAP-25 concentrations, the transition from the inactive state to the active state becomes rate-limiting, causing the probability to plateau.

To test this model, we solved the master equations governing the reaction scheme (Fig. 1F, inset), fit to the measured SNARE assembly probabilities (Figs. 1F & S1), and determined the three unknown rate constants in the model (k_o_, k_c_, and k_b_) (SI Appendix). Two other parameters in the model - the folding rate (k_f_) and unfolding rate (k_u_) of the template complex at the equilibrium force, were fixed to their experimental measurement 3.2 s^−1^ (Jiao et al., 2018). The model predictions generally match the measured probabilities, indicating that our model is consistent with the experimental data. Nonlinear fitting revealed that the template complex transitions from the inactive state to the active state with a rate of 0.05 (± 0.02, S.D.) s^−1^ and an increase in free energy of 6.8 (± 0.3) k_B_T. The relatively slow activation rate limits the SNARE assembly probability at a high SNAP-25 ≥ 0.4 μM. The high energy is likely associated with the conformational changes in the orientation and separation of the two aligned helices required for SNAP-25 binding (Fig. 1B). SNAP-25 quickly associates with the activated template complex, with a bimolecular rate constant of 7 (±5) ×10^7^ M^−1^s^−1^ that approaches the diffusion limit. Thus, the activated template complex nucleates rapid SNARE assembly.

### SNARE crosslinking at the -6 layer significantly attenuates SNARE assembly

In contrast to our studies using Syx-VAMP conjugate crosslinked at X-8, the cryo-EM structure of the template complex was determined using syntaxin-1 and VAMP2 that were crosslinked at X-6 (Fig. 1A) (Stepien et al., 2022). It is unknown how the choice of the crosslinking site affects the structure of the template complex or the pathway or kinetics of chaperoned SNARE assembly. To address this question, we measured the stability and SNAP-25 binding probabilities of the template complex crosslinked at X-6. We found that the X-6 crosslinking supported the efficient formation of the template complex and open syntaxin (Fig. 2A; Table 1), consistent with our previous observations (Jiao et al., 2018). At constant force in the range of 3-9 pN, the template complex reversibly unfolded to open syntaxin in a force-dependent manner (Fig. 2B). The equilibrium force of the template complex crosslinked at X-6 (5.8 pN) is greater than that crosslinked at X-8 (5.1 pN), but the average extension change associated with the former template complex is smaller (4.2 nm vs 5.4 nm). Hidden-Markov modeling of these trajectories revealed the unfolding probabilities and folding and unfolding rates as a function of force (Fig. 2C) (Zhang et al., 2016b). These measurements were fit by a force-dependent model for protein folding (Rebane et al., 2016), yielding an unfolding energy of 3.9 (±0.1, SEM) k_B_T for the template complex crosslinked at X- 6. Thus, X-6 crosslinking destabilizes the template complex by 1.3 k_B_T compared with X-8 crosslinking (Table 1).

**Fig. 2.**
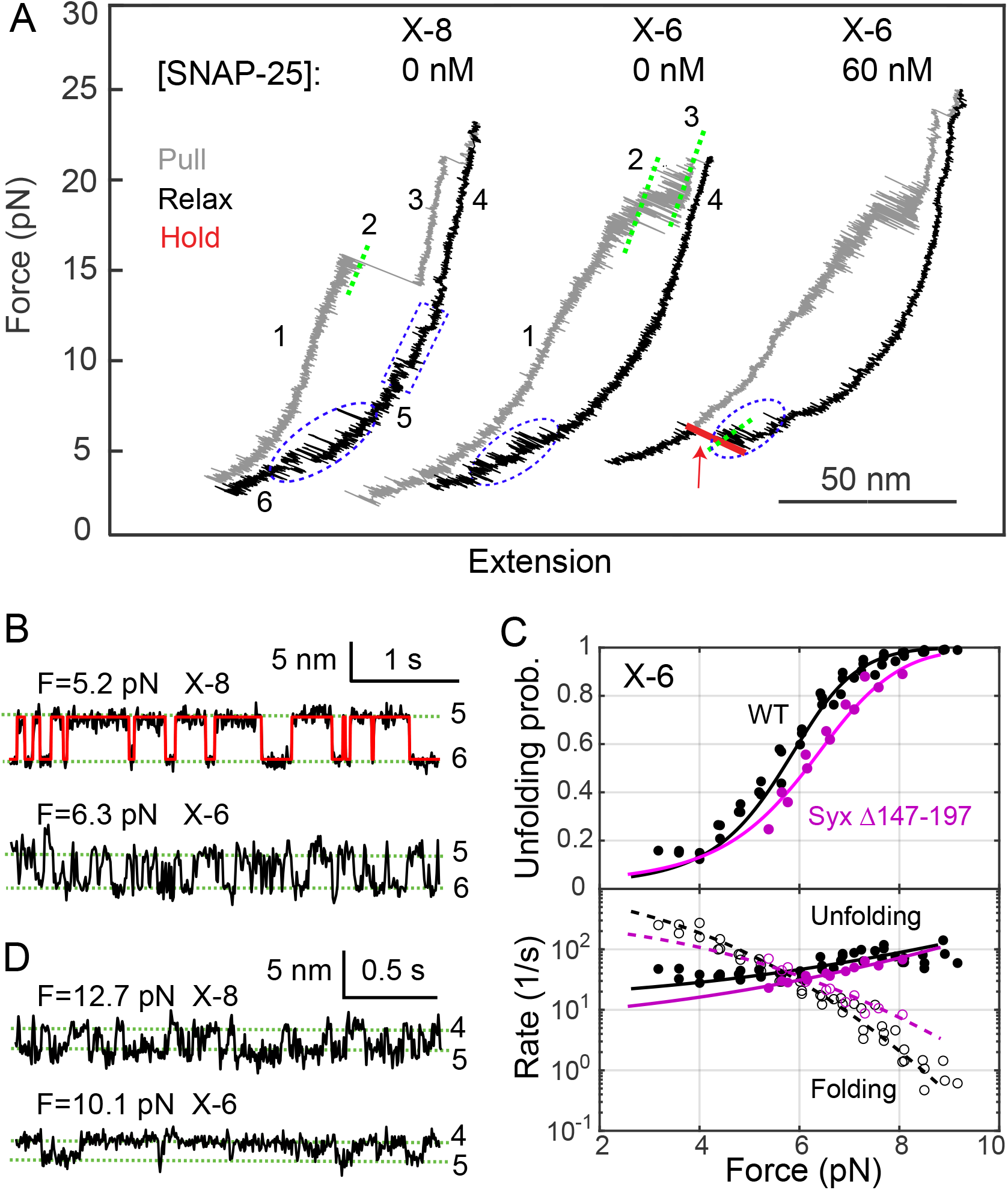
Shifting the crosslinking site from X-8 to X-6 significantly reduces the stability of the template complex and SNAP-25 binding but does not alter the stability of open syntaxin. (A) FECs obtained by manipulating single syntaxin-1-VAMP2 conjugates crosslinked at X-8 or X-6 in the presence of 1 μM Munc18-1. The NTD transition of the SNARE complex (between states 2 and 3) becomes reversible when crosslinked at X-6 (Ma et al., 2015). (B) Extension-time trajectories showing reversible unfolding and refolding of the template complex at constant mean force. The red curve indicates the idealized transition derived from hidden-Markov modeling. (C) Unfolding probabilities (top panel) and folding and unfolding rates (bottom panel) of the template complex crosslinked at X-6 as a function of force. Non-linear model fitting (curves) yielded the unfolding energy and rates at zero force (see Materials and methods). (D) Extension-time trajectories showing folding and unfolding transitions of open syntaxin at a constant force.

Next, we tested SNAP-25 binding to the template complex using the assay described above. The template complex crosslinked at X-6 bound to SNAP-25 with a significantly lower probability than its counterpart crosslinked at X-8 (0.035 vs 0.27) in the presence of 60 nM SNAP-25 (Fig. 2A; Table 1). Increasing the SNAP-25 concentration to 120 nM led to binding probabilities of 0.086 for X-6 (N=35) and of 0.32 for X-8 (N=34). Therefore, although crosslinking at X-6 maintains the template complex, it significantly reduces SNAP-25 binding compared to crosslinking at X-8.

### Syntaxin linker region is required to activate the template complex for SNARE assembly

Inspired by the cryo-EM structure of the template complex, we examined the role of the syntaxin linker region (a.a. 147-197) in Munc18-1-chaperoned SNARE assembly. To this end, we tested the effects of four modifications in the syntaxin linker on SNAP-25 binding and the stabilities of the template complex and open syntaxin. The syntaxin linker is partially folded in both closed syntaxin and the template complex and likely disordered in the fully assembled SNARE complex (Figs. 3A & 1B) (Misura et al., 2000; Stepien et al., 2022). We and others have previously shown that the syntaxin linker stabilizes closed syntaxin, consistent with its structure (Burkhardt et al., 2008; Jiao et al., 2018). Surprisingly, we found that truncation of the entire linker region (Syx Δ147-197) does not significantly alter the stabilities of both template complex and open syntaxin, regardless of the crosslink site (X-8 or X-6) (Fig. 2C; Fig. 3B, FEC #1 & #4; Fig. 3C; Table 1). This surprising finding suggests that the syntaxin linker region and likely the whole N-terminal regions of SNARE motifs are highly flexible. However, the syntaxin truncation impairs SNAP-25 binding, with no binding observed for X-8 and a low probability of 0.029 for X-6. Consistent with these results, two mutations M183A and D184P in the syntaxin linker region (Fig. 1A, inset), did not affect the stabilities of the template complex and open syntaxin (Fig. S2; Table 1). Intriguingly, both mutations were shown to abolish the association between Munc18-1 and the Syx-VAMP conjugate (Stepien et al., 2022). The apparent discrepancy will be discussed later. Taken together, these observations suggest that the syntaxin linker region minimally affects the stability of the inactive template complex, specifically stabilizes the activated template complex, and is essential for Munc18-1-chaperoned SNARE assembly.

**Fig. 3.**
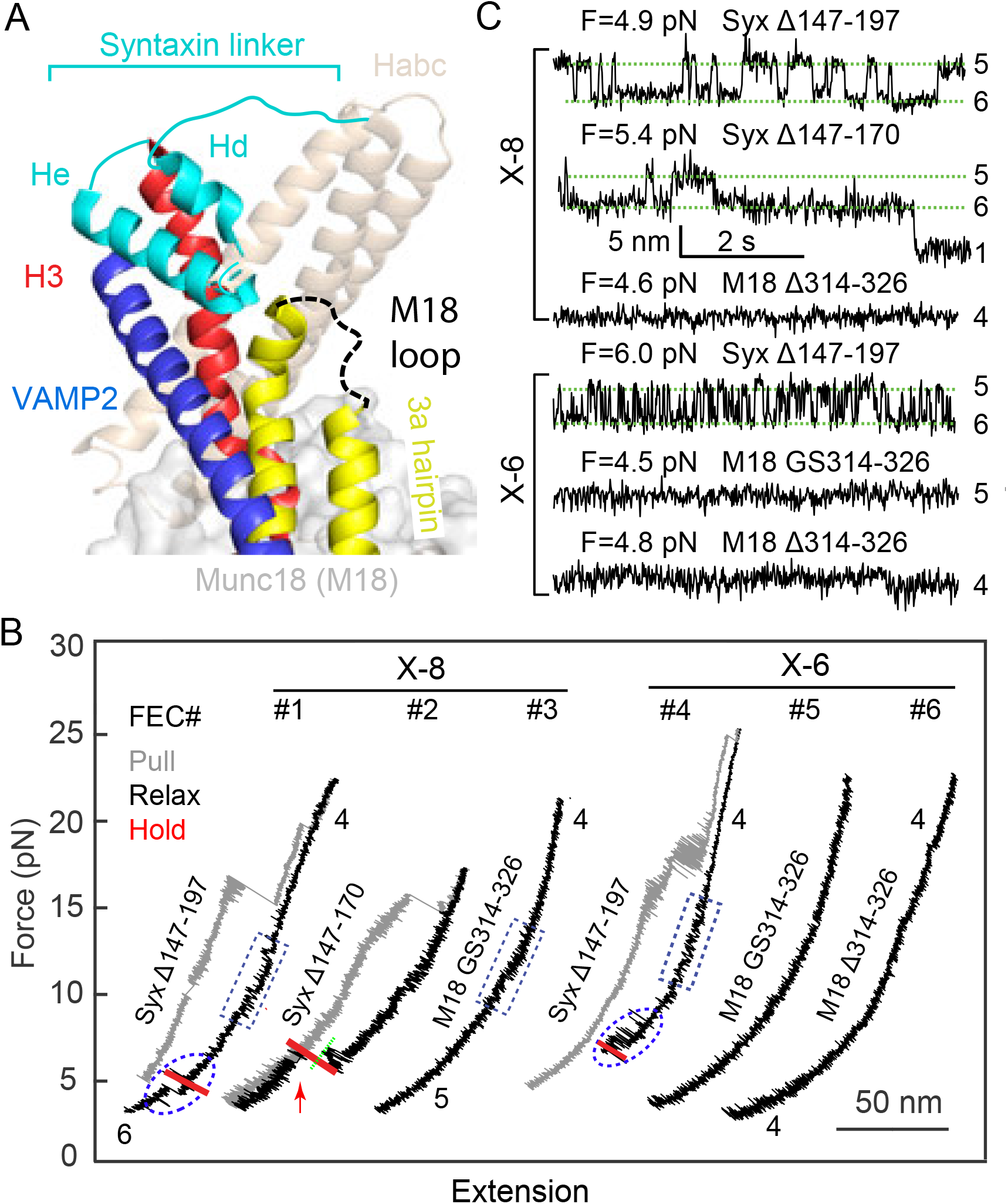
The syntaxin linker region (a.a. 147-197) and the Munc18-1 loop in the 3a helical hairpin domain (a.a. 314-326) are required for chaperoned SNARE assembly with different mechanisms. (A) Positions and conformations of the two regions in the template complex. (B) FECs associated with truncated syntaxin-1 (Syx Δ147-197) or Munc18-1 (M18 Δ314-326), or Munc18-1 with the loop replaced by the glycine- and serine-rich sequence (M18 GS314-326) in the presence of 1 μM Munc18-1 and 60 nM SNAP-25 in the solution. (C) Extension-time trajectories at constant mean force for Syx-VAMP conjugates crosslinked at X-8 or X-6.

To pinpoint the syntaxin linker region required for the chaperoned SNARE assembly, we tested the template complex crosslinked at X-8 with partial linker truncation (Syx Δ147-170). Interestingly, the truncation barely affects the template complex and SNAP-25 binding (Fig. 3B, FEC #2; Fig. 3C; Table 1). Combining the results from both partial and full linker truncations, we conclude that the C-terminal region of the syntaxin linker (a.a. 171-197), including the He helix seen in the cryo-EM structure of the template complex (Fig. 3A), is essential for activation of the template complex and SNAP-25 binding, while the N-terminal region (a.a. 147-170), including the Hd helix, may be dispensable at least in our assay.

### Munc18-1 3a hairpin loop is essential for SNARE assembly

The tip of the Munc18-1 3a hairpin (a.a. 314-334) contains a loop (a.a. 314-326) that is disordered in all published Munc18-1 structures (Fig. 3A) (Misura et al., 2000; Stepien et al., 2022). The function of this loop is unclear. In closed syntaxin, it furls back as part of the tip and appears to associate with the 3a helices to stabilize the furled conformation. Thus, mutations around or in the loop, such as P335A and D326K, destabilize closed conformation, stabilize the template complex and VAMP2 binding, and promote SNARE assembly and neurotransmission (Andre et al., 2020; Jiao et al., 2018; Munch et al., 2016; Parisotto et al., 2014; Sitarska et al., 2017). Yet, the loop does not appear to bind any SNAREs or other regions of Munc18-1 in the cryo-EM structure of the template complex (Stepien et al., 2022). Thus, the specific role of this loop in template complex formation and SNARE assembly needs further investigation.

To examine the function of the loop, we truncated the loop (M18 Δ314-326) or replaced it with a glycine- and serine-rich sequence (M18 GS314-326) and then measured the template complex stability and SNARE assembly. Surprisingly, we found that the two modifications abolished both the template complex and SNAP-25 binding (Fig. 3B, FEC #3, #5, #6; Fig. 3C; Table 1). Thus, the loop is essential for template complex formation and Munc18-1-chaperoned SNARE assembly. In addition, while the loop substitution did not affect open syntaxin (Fig. 3B, FEC #3), the loop truncation abolished it (FEC #6; Table 1). Both observations are consistent with the structural model of open syntaxin (Eisemann et al., 2020; Jiao et al., 2018; Zhang and Hughson, 2021). In the structures of the template complex, and likely open syntaxin, the syntaxin SNARE motif contacts Munc18-1 3a hairpin primarily at a helical region (a.a. 335-339) adjacent to the loop. The loop truncation likely pulls the helical region away from the SNARE motif to destabilize the interaction (Munch et al., 2016), but the loop substitution does not. Therefore, the loop directly stabilizes the template complex, likely by interacting with VAMP2 or syntaxin-1, rather than destabilizing open syntaxin.

### Template complexes mediate specific SNARE associations

To examine whether template complexes specifically bind Qbc-SNAREs, we tested the binding of SNAP-25 and SNAP-23 to two template complexes Munc18-1:Syntaxin-1:VAMP2 and Munc18-3:Syntaxin-4:VAMP2 (Jiao et al., 2018). All preassembled SNARE complexes show characteristic stepwise SNARE unfolding behavior (Fig. 4A, grey FECs) (Zorman et al., 2014). This observation indicates that, in the absence of Munc18, syntaxin-1 and syntaxin-4 nonspecifically pair with either SNAP-23 or SNAP-25 to form SNARE complexes in the presence of VAMP2, consistent with previous results (Bajohrs et al., 2005). In addition, the SNARE complexes containing SNAP-25 have similar CTD zippering kinetics and energies as those containing SNAP-23 (Fig. 4B), with CTD zippering energies of 27 (±3, SD) k_B_T for Syntaxin-4:VAMP2:SNAP-25 and 24 (±2) k_B_T for Syntaxin-4:VAMP2:SNAP- 23. This result is not surprising, because SANP-25 and SNAP-23 share ∼ 60% identical amino acids and only differ three residues at -6 and -5 hydrophobic layers and one residue at the +8 layer. These comparisons suggest that the different kinetics of exocytosis mediated by these SNAREs may be controlled by regulatory proteins (Kadkova et al., 2019; Thurmond and Gaisano, 2020).

**Fig. 4.**
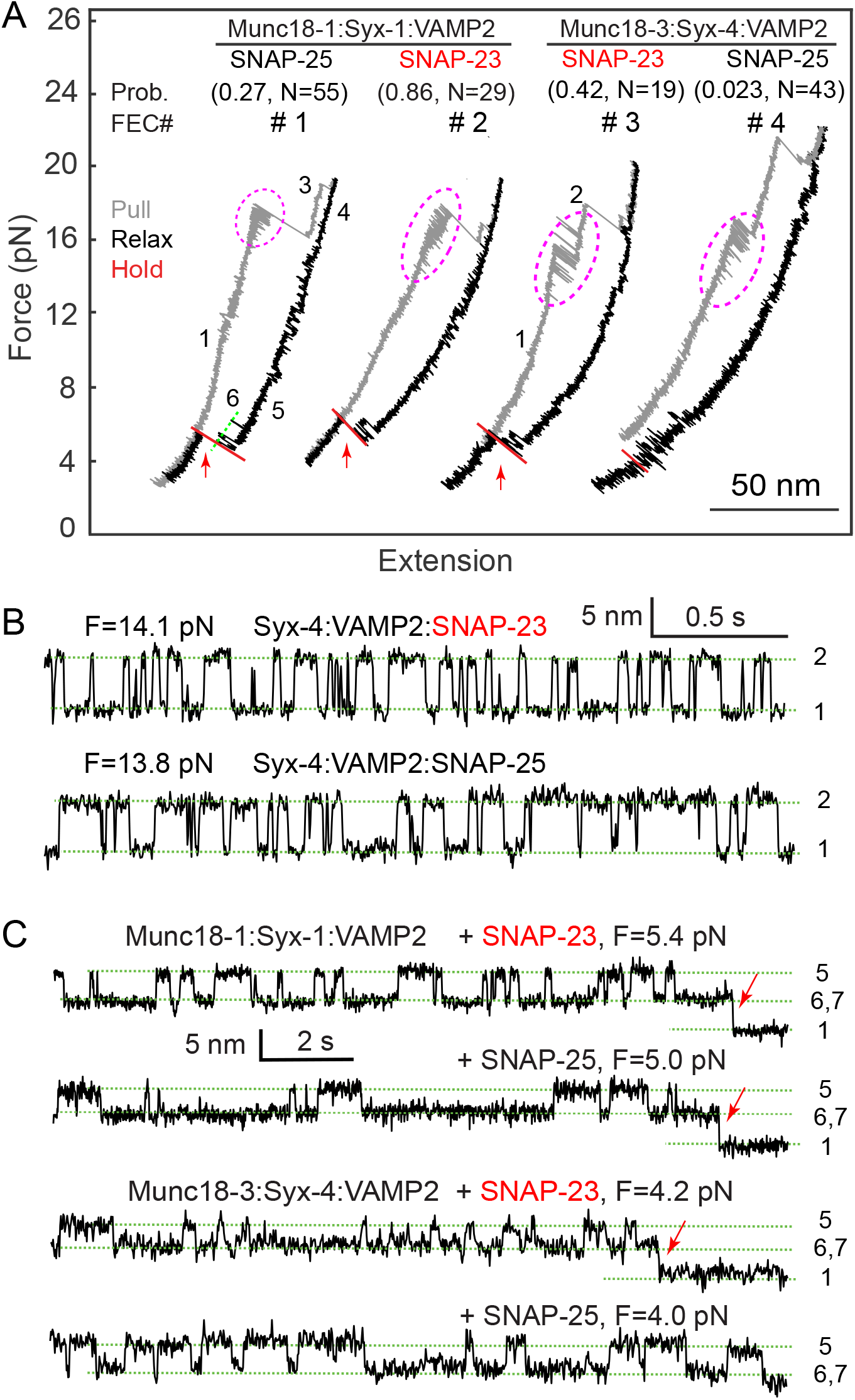
The template complexes proofread Qbc-SNARE binding. (A) FECs showing specific Qbc SNARE (SNAP-25 or SNAP-23) binding to its cognate template complex containing either Munc18-1 or Munc18-3. The Qbc-SNARE binding probabilities and associated total numbers of tests are shown in pairs in parenthesis. Reversible CTD transitions are marked by magenta dashed ovals. (B) Extension-time trajectories at constant mean force showing folding and unfolding transitions of template complexes and Qbc-SNARE binding (indicated by red arrows).

Indeed, Munc18 proteins regulate the speed or specificity of SNARE assembly. We quantified the specificity of Qbc-SNARE binding based on their binding probabilities. SNAP-23 bound to the synaptic template complex with an even higher probability than SNAP-25 (0.86 vs 0.27; Fig. 4, FEC #1 & #2), which is consistent with the finding that SNAP-23 supports neurotransmitter release in SNAP-25 knockout neurons (Delgado-Martinez et al., 2007). In contrast, SNAP-23 is strongly favored by Munc18-3:Syntaxin-4:VAMP2 compared with SNAP-25, as is demonstrated by their different binding probabilities (0.42 vs 0.023) (Fig. 4, FEC #3 & #4). Increasing the SNAP-25 concentration to 180 nM slightly increased the binding probability to 0.09 (N=34). Thus, both template complexes confer specificity for binding of Qbc SNAREs, consistent with the specialized role of different SNARE/SM fusion machines in different membrane trafficking pathways (Koike and Jahn, 2022; Shen et al., 2007). Considering the close CTD zippering energies of the resultant SNARE complexes, we conclude that Munc18 proteins proofread Qbc-SNARE association with template complexes primarily based on their NTDs, as the splayed CTDs of Qa- and R-SNAREs on the SM proteins cannot initiate the zippering process.

## Discussion

Using high-resolution optical tweezers, we find that the synaptic template complex undergoes a conformational transition to a high-energy state for SNAP-25 binding. The activated state requires a structural rearrangement – likely folding of ∼ 30 amino acids in the linker region of syntaxin immediately N-terminal to its SNARE motif, as its truncation barely affects the stability of the template complex but impairs SNAP-25 binding. These findings corroborate the recent cryo-EM structure of the template complex crosslinked at X-6 (Stepien et al., 2022) and previous observation that the 20 amino acids N-terminal to the SNARE motif of the yeast Qa-SNARE Vam3 are essential for SNARE assembly chaperoned by the cognate SM protein Vps33 (Song and Wickner, 2017). Thus, the linker regions N-terminal to the SNARE motifs of Qa-SNAREs may be generally required for SM-chaperoned SNARE assembly. We also observe that SNARE crosslinking at X-6 significantly reduces the activity of the template complex to bind SNAP-25 compared to X-8 (Table 1). Thus, crosslinking at X-6 may alter the conformation of the native template complex, for example, causing detachment of the VAMP2 NTD from the Munc18-1 surface. Supporting this view, we found that the loop in Munc18-1 3a hairpin is required for template complex formation, which implies that the VAMP2 NTD interacts with the loop region in the native template complex. This finding is consistent with the crystal structure of Vps33 complexed with its cognate R-SNARE Nyv1, in which its NTD makes extensive contact with the Vps33 3a hairpin (Baker et al., 2015). Likewise, crosslinking at X-6 may stabilize the Hd helix seen in the cryo-EM structure. Nevertheless, the resolution of the Hd helix is too low to resolve its amino acid sequence, implying its high conformational flexibility. Our data indicate that the entire syntaxin linker region is highly dynamic. Particularly, the Hd helix is dispensable for the formation of the template complex and SNAP-25 binding. Finally, the cryo-EM study also revealed an alternative structure suggested to represent an intermediate transitioning from the closed syntaxin to the template complex (Stepien et al., 2022). This structure exhibits a slightly different conformation compared with the mature conformation described above; in particular, the two NTDs lie closer to the underlying Munc18-1 surface, confirming the conformational flexibility of the template complex. Taken together, while our data are generally consistent with the cryo-EM structure, our findings demonstrate an even more dynamic structure of the template complex that is essential for chaperoned SNARE assembly and susceptible to perturbation by SNARE crosslinking.

Our single-molecule optical tweezers approach allowed us to resolve at least seven states involved in Munc18-1-chaperoned SNARE assembly, and to measure their stabilities and transition kinetics (Fig. 1D). Accordingly, we report the stability of the template complex as the energy difference between the template complex and open syntaxin. In contrast, Stepien et al. used mass photometry to evaluate the binding between Munc18-1 and the Syx-VAMP conjugate crosslinked at X-6 (Stepien et al., 2022). This single-molecule method could only revolve free and SNARE-bound Munc18-1. Based on the reaction scheme shown in Fig. 1D, the dissociation energy includes the free energies of the template complex, the open syntaxin, and the dissociation energy between Munc18-1 and the N-terminal domains of syntaxin (Fig. 1D, state 4) (Burkhardt et al., 2008). The comparison suggests that the two single-molecule approaches measured the stabilities of the template complex relative to different reference states. While mass photometry showed that the two mutations in the syntaxin linker M183A and D184P abolished the association between Munc18-1 and the Syx-VAMP conjugate (Stepien et al., 2022), our assay revealed that both mutations barely affected the stability of the template complex relative to open syntaxin. The comparison suggests that these mutations likely destabilize other states leading to Munc18-1 dissociation.

Our data suggest an intriguing interplay between the specificity and stability of chaperoned assembly of SNAREs or protein complexes in general. Given a mixture of SNAREs in the absence of chaperones, the specificity of their pairing under an equilibrium condition is determined by the Boltzmann distribution or ultimately by the relative stabilities of the SNARE complexes. However, in the presence of chaperones, the specificity is no longer determined by the relative stabilities and the resultant SNARE complexes are kinetically trapped given their extremely long lifetimes. Consistent with this view, SM proteins proofread the association of Qbc-SNAREs based on their NTDs, instead of the entire SNARE motifs. We believe that this view may help further dissect the roles of different SM and SNARE proteins in membrane fusion in the same trafficking pathways, such as insulin release (Thurmond and Gaisano, 2020).

Due to its essential role in SNARE assembly, the dynamic template complex, especially the activated template complex, may be regulated by other proteins to control membrane fusion. We have previously shown that Munc13-1 binds to the template complex to enhance SNARE assembly and that Munc18-1 phosphorylation can modulate SNARE assembly (Jiao et al., 2018; Shu et al., 2020; Zhang and Hughson, 2021). Future work is required to pinpoint the role of the activated template in regulated SNARE assembly and determine the structure of the template complex in the presence of X-8 crosslinking or the absence of any SNARE crosslinking.

## Materials and Methods

### Protein Constructs and Purification

The amino acid sequences corresponding to WT SNARE and Munc18-1 and their purification were described elsewhere in detail (Jiao et al., 2018). Briefly, the genes containing the cytoplasmic domains of rat syntaxin-1A (a.a. 1-265 with mutations C145S and R198C or L205C), syntaxin-4 (a.a. 1-273 with Q194C), VAMP2 (a.a. 1-96 with N29C or Q36C), and rat Munc18-1 were cloned into the pET-SUMO vector encoding 6xHis-tag followed by a SUMO tag at the N termini. The coding sequence for rat Munc18-3 was cloned into pET-15a (Novagen, TX) and codon-optimized for protein expression in bacteria. The genes for the mutant proteins were derived by PCR mutagenesis. Particularly, Munc18-1 mutant M18 GS314-326 had a loop replaced by GSGGRGNGGSAGS. The full-length cysteine-freer SNAP-25B (C85S, C88S, C90S, C92S) and SNAP-23 (C79S, C80S, C83S, C85S, C87S) were cloned into pET-15b vector encoding 6xHis-tag at the N terminus. These proteins were expressed in BL21 E. coli cells and purified using Ni-NTA-agarose beads. Proteins were eluted with 300 mM imidazole and exchanged to buffer containing 25 mM HEPES (pH 7.4), 140 mM KCl, and 2 mM tris(2- carboxyethyl)phosphine (TCEP). Syntaxin-1A and syntaxin-4 were then biotinylated at their C-terminal Avi-tags with the biotin ligase BirA as previously described.

### SNARE Complex Formation and Crosslinking to DNA handle

Ternary SNARE complexes were prepared and cross-linked with DNA handles as was previously described (2, 4, 5). Syntaxin- 1A, SNAP-25B, and VAMP2 were mixed at a molar ratio of 0.8:1:1.2, incubated at 4 °C, and purified using the 6xHis-tag on SNAP-25B and Ni-NTA-agarose. The eluted SNARE complexes were cross-linked with DTDP (2,2’-dithiodipyridine disulfide) treated DNA handles with a molar ratio of 50:1 in 100 mM phosphate buffer, 500 mM NaCl, pH 8.5. Other preassembled SNARE complexes were similarly prepared.

### High-resolution dual-trap optical tweezers

The optical tweezers were home-built as described elsewhere (Moffitt et al., 2006). Briefly, a 1064 nm laser beam is expanded, collimated, and split into two orthogonally polarized beams, one of which is reflected by a mirror attached to a nano- positioning stage (Mad-city Labs, WI). The two beams are then combined, expanded again, and are then focused by a water-immersed 60x objective with a 1.2 numerical aperture (Olympus, PA) to form two optical traps in the sample plane in the central channel of a home-built microfluidic flow chamber. One of the two traps is stationary; the other trap can be moved using the nano- positioning stage. The outgoing laser beams are collimated by a second water-immersed objective, split again by polarization, and projected onto two position-sensitive detectors (Pacific Silicone Sensor, CA). Displacements of the trapped beads are detected by back-focal plane interferometry. Optical tweezers are remotely operated through a computer interface written in LabVIEW (National Instruments, TX). The force constants of two optical traps are determined by the Brownian motion of the trapped beads before each experiment.

### Single-molecule protein folding experiment

An aliquot of the crosslinked protein-DNA sample was incubated with 1 μL anti-digoxigenin coated polystyrene beads 2.17 μm in diameter (Spherotech, IL), diluted in 1 mL phosphate-buffered saline (PBS), and injected into the top channel of a microfluidic chamber (Jiao et al., 2017). Streptavidin-coated polystyrene beads of 1.86 μm were injected into the bottom channel. Both top and bottom channels were connected to a central channel by capillary tubes, where both kinds of beads were trapped. A single SNARE complex was tethered between two beads by bringing them close. Data were recorded at 20 kHz, mean-filtered to 10 kHz, and stored on a hard disc. The single-molecule experiment was conducted in PBS at 23 (±1) °C. An oxygen scavenging system was added to prevent potential protein photo- damage by optical traps. The single protein-DNA tether was pulled or relaxed by increasing or decreasing trap separation at a speed of 10 nm/s.

### Data analysis

Our methods were described in detail elsewhere (Jiao et al., 2017; Rebane et al., 2016; Zhang et al., 2016b). Briefly, the extension trajectories were analyzed by two-state hidden-Markov modeling (HMM), which yielded the probability, extension, force, lifetime, and transition rates for each state (Zhang et al., 2016b). To relate the experimental measurements to the conformations and energy (or the energy landscape) of different SNARE states at zero force, we constructed structural models for these states based on crystal structures of the SNARE four-helix bundle and the template complex (Rebane et al., 2016). These states were characterized by the contour lengths of the unfolded polypeptides and free energy, which were chosen as fitting parameters. The extension and energy of the whole tethered dumbbell, including the DNA handle, were calculated using the Marko-Siggia formula (Marko and Siggia, 1995). Then, we computed the probability of each state based on the Boltzmann distribution and transition rates based on the Kramers’ equation. Finally, we fit the calculated state extensions, forces, probabilities, and transition rates to the corresponding experimental measurements using the nonlinear least-squares fitting, which revealed the conformations and energies of different SNARE folding states as best-fit parameters.

## Acknowledgements

We thank Fred Hughson and Jose Rizo for reading our manuscript and constructive discussion, and Avinash Kumar for technical assistance. This work was supported by NIH grants R35 GM131714 to Y. Z. Y. Z. and J. Y. designed the experiments, J. Y., H. J., Y. L. and Y. G. performed the experiments, Y. Z., J. Y. and H. J. analyzed the data, and wrote the paper. The authors declare no interest of conflict.

## Supporting Information for

The SI Appendix contains supplemental text and two supplemental figures.

### Kinetic model of the conformational transitions of the template complex and SNAP-25 binding

Suppose the template complex undergoes reversible and sequential conformational transitions among the unfolded state, the folded but inactive closed state, and the folded and active open state to which SNAP-25 can bind, as shown in the inset in Fig. 1F with indicated transition rates. Then the probabilities to observe the unfolded template complex *p*_*u*_, the closed template complex *p*_*c*_, the open template complex *p*_*o*_, and the fully assembled SNARE complex *p*_*s*_ at any time *t* are determined by the Master equations:

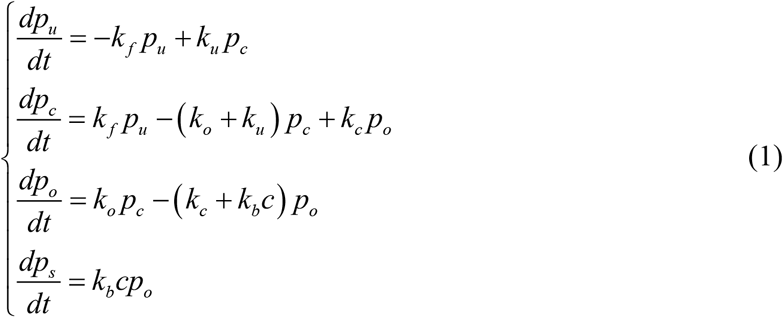

where *c* is the concentration of SNAP-25 in the solution. Defining the probability vector

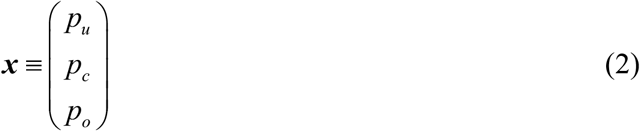

and the rate matrix

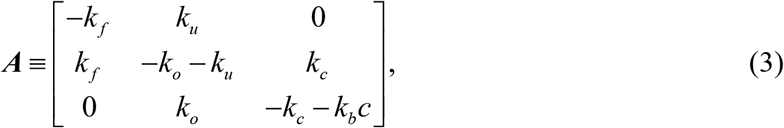

We can rewrite Eq. (1) as

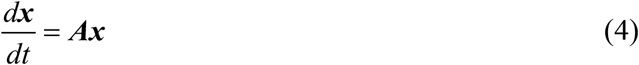

and

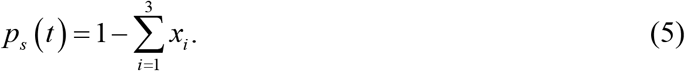

Suppose the rate matrix has the eigenvalues *λ*_*i*_, *i* = 1, …, *n* and the corresponding eigenvectors ***v***_*i*_ such that

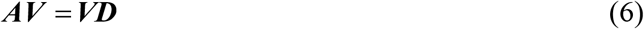

where ***V*** is a matrix formed by all the eigenvectors

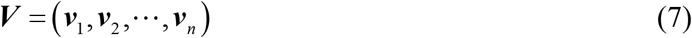

and ***D*** is a diagonal matrix

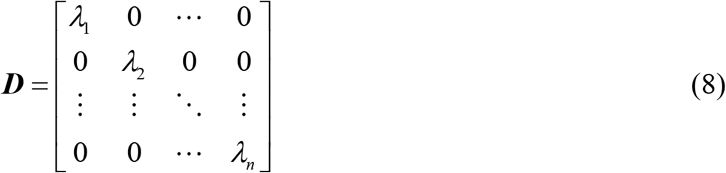

The solution to Eq. (4) is

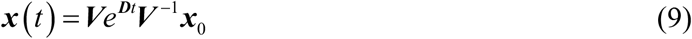

Where ***x***_0_ is a constant vector whose elements represent the initial state probabilities at time zero.

Given model parameters, we numerically computed the state probabilities according to Eqs. (9) and (5) using MATLAB. These calculations were fit to the experimental measurements to derive the rate constants in the model as best-fit parameters. In our SNAP-25 binding experiments, we held the Syx-VAMP conjugate at a constant mean force close to the equilibrium force of the template complex. In this case, the template complex folded and unfolded with equal rates, which were previously measured, i.e., *k* _*f*_ = *k*_*u*_ = 3.2 s^−1^. Correspondingly, the initial state probabilities were chosen as ***x***_0_ = (0.5, 0.5, 0). Thus, our model has three rate constants as fitting parameters. The SNAP-25 binding probabilities were measured at variable SNAP-25 concentration with the fixed time (t=120 s, Fig. 1F) or variable time with fixed SNAP-25 concentration (60 nM, Fig. S1). These probabilities were fit by the model predictions to determine the three rate constants (Fig. 1F). The MATLAB scripts used for these calculations are available upon request.

**Fig. S1.**
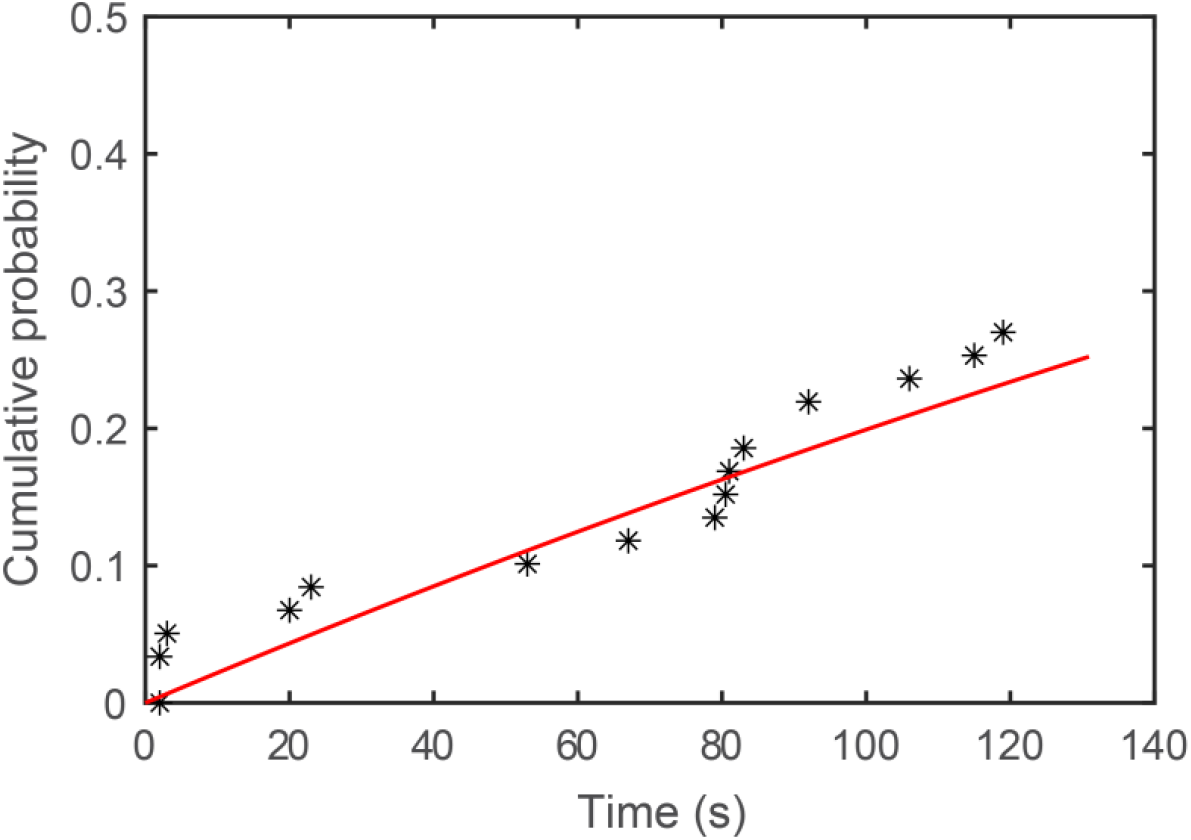
Cumulative probability calculated from the measured time for SNAP-25 binding in the presence of 60 nM SNAP-25 in the solution (symbols) and its best model fit (red curve).

**Fig. S2.**
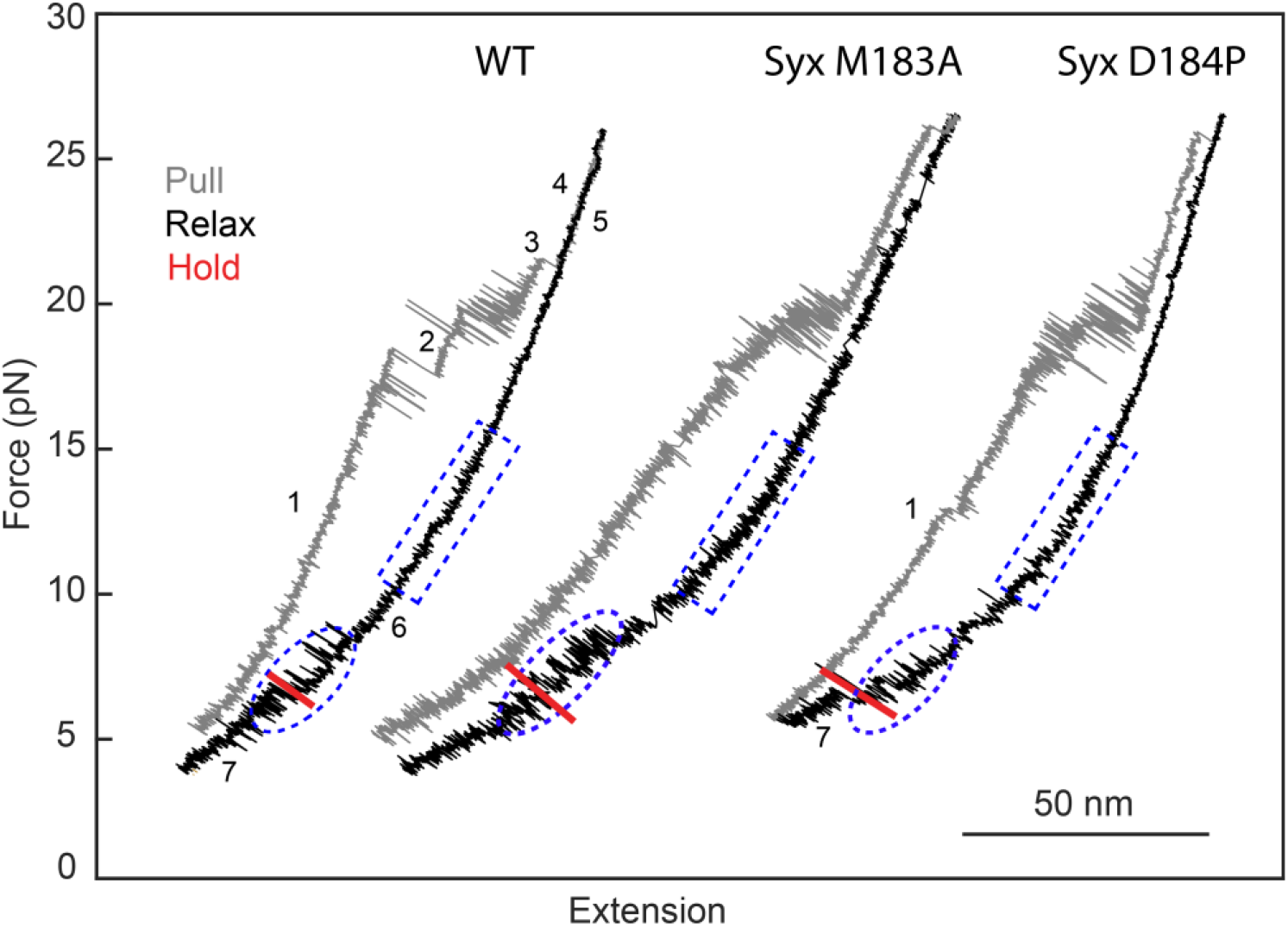
FECs associated with syntaxin-1 with point mutation M183A or D184P in the linker in the presence of 1 μM Munc18-1 and 60 nM SNAP-25 in the solution. Single pre-assembled SNARE complexes were first pulled to high forces to detect their stepwise unfolding (grey FECs) and then relaxed to low forces to detect reversible refolding of open syntaxin (blue dashed rectangles) and template complexes (blue dashed ovals). Finally, the complexes were held around the equilibrium of the template complex transitions to detect SNAP-25 binding (red regions). The states associated with different FEC regions are labeled as in Fig. 1D.

## References

Andre, T., Classen, J., Brenner, P., Betts, M.J., Dorr, B., Kreye, S., Zuidinga, B., Meijer, M., Russell, R.B., Verhage, M., et al. (2020). The interaction of Munc18-1 Helix 11 and 12 with the central region of the VAMP2 SNARE motif is essential for SNARE templating and synaptic transmission. Eneuro 7, ENEURO.0278-0220.2020.

Bajohrs, M., Darios, F., Peak-Chew, S.Y., and Davletov, B. (2005). Promiscuous interaction of SNAP-25 with all plasma membrane syntaxins in a neuroendocrine cell. Biochem J 392, 283–289.

Baker, R.W., and Hughson, F.M. (2016). Chaperoning SNARE assembly and disassembly. Nat Rev Mol Cell Biol 17, 465–479.

Baker, R.W., Jeffrey, P.D., Zick, M., Phillips, B.P., Wickner, W.T., and Hughson, F.M. (2015). A direct role for the Sec1/Munc18-family protein Vps33 as a template for SNARE assembly. Science 349, 1111–1114.

Brunger, A.T. (2005). Structure and function of SNARE and SNARE-interacting proteins. Q Rev Biophys 38, 1–47.

Burkhardt, P., Hattendorf, D.A., Weis, W.I., and Fasshauer, D. (2008). Munc18a controls SNARE assembly through its interaction with the syntaxin N-peptide. EMBO J 27, 923–933.

Cecconi, C., Shank, E.A., Bustamante, C., and Marqusee, S. (2005). Direct observation of the three-state folding of a single protein molecule. Science 309, 2057–2060.

Choi, U.B., Zhao, M.L., White, K.I., Pfuetzner, R.A., Esquivies, L., Zhou, Q.J., and Brunger, A.T. (2018). NSF-mediated disassembly of on-and off-pathway SNARE complexes and inhibition by complexin. Elife 7, e36497.

Delgado-Martinez, I., Nehring, R.B., and Sorensen, J.B. (2007). Differential abilities of SNAP-25 homologs to support neuronal function. J Neurosci 27, 9380–9391.

Eisemann, T.J., Allen, F., Lau, K., Shimamura, G.R., Jeffrey, P.D., and Hughson, F.M. (2020). The Sec1/Munc18 protein Vps45 holds the Qa-SNARE Tlg2 in an open conformation. Elife 9, e60724.

Fasshauer, D., Sutton, R.B., Brunger, A.T., and Jahn, R. (1998). Conserved structural features of the synaptic fusion complex: SNARE proteins reclassified as Q-and R-SNAREs. Proc Natl Acad Sci USA 95, 15781–15786.

Gao, Y., Zorman, S., Gundersen, G., Xi, Z.Q., Ma, L., Sirinakis, G., Rothman, J.E., and Zhang, Y.L. (2012). Single reconstituted neuronal SNARE complexes zipper in three distinct stages. Science 337, 1340–1343.

Jiao, J., He, M., Port, S.A., Baker, R.W., Xu, Y., Qu, H., Xiong, Y., Wang, Y., Jin, H., Eisemann, T.J., et al. (2018). Munc18-1 catalyzes neuronal SNARE assembly by templating SNARE association. Elife 7, e41771.

Jiao, J.Y., Rebane, A.A., Ma, L., and Zhang, Y.L. (2017). Single-molecule protein folding experiments using high-resolution optical tweezers. Methods Mol Biol 1486, 357–390.

Kadkova, A., Radecke, J., and Sorensen, J.B. (2019). The SNAP-25 protein family. Neuroscience 420, 50–71.

Koike, S., and Jahn, R. (2022). SNARE proteins: zip codes in vesicle targeting? Biochem J 479, 273–288.

Ma, C., Su, L.J., Seven, A.B., Xu, Y.B., and Rizo, J. (2013). Reconstitution of the vital functions of Munc18 and Munc13 in neurotransmitter release. Science 339, 421–425.

Ma, L., Rebane, A.A., Yang, G., Xi, Z., Kang, Y., Gao, Y., and Zhang, Y.L. (2015). Munc18-1-regulated stage-wise SNARE assembly underlying synaptic exocytosis. Elife 4, e09580.

Marko, J.F., and Siggia, E.D. (1995). Stretching DNA. Macromolecules 28, 8759–8770.

Misura, K.M.S., Scheller, R.H., and Weis, W.I. (2000). Three-dimensional structure of the neuronal-Sec1-syntaxin 1a complex. Nature 404, 355–362.

Moffitt, J.R., Chemla, Y.R., Izhaky, D., and Bustamante, C. (2006). Differential detection of dual traps improves the spatial resolution of optical tweezers. Proc Natl Acad Sci USA 103, 9006–9011.

Munch, A.S., Kedar, G.H., van Weering, J.R.T., Vazquez-Sanchez, S., He, E.Q., Andre, T., Braun, T., Sollner, T.H., Verhage, M., and Sorensen, J.B. (2016). Extension of Helix 12 in Munc18-1 induces vesicle priming. J Neurosci 36, 6881–6891.

Parisotto, D., Pfau, M., Scheutzow, A., Wild, K., Mayer, M.P., Malsam, J., Sinning, I., and Sollner, T.H. (2014). An extended helical conformation in domain 3a of Munc18-1 provides a template for SNARE (soluble N-ethylmaleimidesensitive factor attachment protein receptor) complex assembly. J Biol Chem 289, 9639–9650.

Pobbati, A.V., Stein, A., and Fasshauer, D. (2006). N-to C-terminal SNARE complex assembly promotes rapid membrane fusion. Science 313, 673–676.

Rebane, A.A., Ma, L., and Zhang, Y.L. (2016). Structure-based derivation of protein folding intermediates and energies from optical tweezers. Biophys J 110, 441–454.

Rebane, A.A., Wang, B., Ma, L., Qu, H., Coleman, J., Krishnakumar, S.S., Rothman, J.E., and Zhang, Y.L. (2018). Two disease-causing SNAP-25B mutations selectively impair SNARE C-terminal assembly. J Mol Biol 430, 479–490.

Rizo, J. (2022). Molecular mechanisms underlying neurotransmitter release. Annu Rev Biophys 51, 377–408.

Shen, J.S., Tareste, D.C., Paumet, F., Rothman, J.E., and Melia, T.J. (2007). Selective activation of cognate SNAREpins by Sec1/Munc18 proteins. Cell 128, 183–195.

Shu, T., Jin, H., Rothman, J.E., and Zhang, Y.L. (2020). Munc13-1 MUN domain and Munc18-1 cooperatively chaperone SNARE assembly through a tetrameric complex. Proc Natl Acad Sci USA 117, 1036–1041.

Sitarska, E., Xu, J.J., Park, S., Liu, X.X., Quade, B., Stepien, K., Sugita, K., Brautigam, C.A., Sugita, S., and Rizo, J. (2017). Autoinhibition of Munc18-1 modulates synaptobrevin binding and helps to enable Munc13-dependent regulation of membrane fusion. Elife 6, e24278.

Sollner, T., Whiteheart, S.W., Brunner, M., Erdjument-Bromage, H., Geromanos, S., Tempst, P., and Rothman, J.E. (1993). SNAP receptors implicated in vesicle targeting and fusion. Nature 362, 318–324.

Song, H., and Wickner, W. (2017). A short region upstream of the yeast vacuolar Qa-SNARE heptad-repeats promotes membrane fusion through enhanced SNARE complex assembly. Mol Biol Cell 28, 2282–2289.

Stamberger, H., Nikanorova, M., Willemsen, M.H., Accorsi, P., Angriman, M., Baier, H., Benkel-Herrenbrueck, I., Benoit, V., Budetta, M., Caliebe, A., et al. (2016). STXBP1 encephalopathy: A neurodevelopmental disorder including epilepsy. Neurology 86, 954–962.

Stepien, K.P., Prinslow, E.A., and Rizo, J. (2019). Munc18-1 is crucial to overcome the inhibition of synaptic vesicle fusion by alpha SNAP. Nat Commun 10, 4326.

Stepien, K.P., Xu, J., Zhang, X., Bai, X.C., and Rizo, J. (2022). SNARE assembly enlightened by cryo-EM structures of a synaptobrevin-Munc18-1-syntaxin-1 complex. Sci Adv 8, eabo5272.

Sudhof, T.C., and Rothman, J.E. (2009). Membrane fusion: grappling with SNARE and SM proteins. Science 323, 474–477.

Sutton, R.B., Fasshauer, D., Jahn, R., and Brunger, A.T. (1998). Crystal structure of a SNARE complex involved in synaptic exocytosis at 2.4 angstrom resolution. Nature 395, 347–353.

Thurmond, D.C., and Gaisano, H.Y. (2020). Recent insights into beta-cell exocytosis in type 2 diabetes. J Mol Biol 432, 1310–1325.

Verhage, M., Maia, A.S., Plomp, J.J., Brussaard, A.B., Heeroma, J.H., Vermeer, H., Toonen, R.F., Hammer, R.E., van den Berg, T.K., Missler, M., et al. (2000). Synaptic assembly of the brain in the absence of neurotransmitter secretion. Science 287, 864–869.

Weninger, K., Bowen, M.E., Choi, U.B., Chu, S., and Brunger, A.T. (2008). Accessory proteins stabilize the acceptor complex for synaptobrevin, the 1 : 1 syntaxin/SNAP-25 complex. Structure 16, 308–320.

Yang, B., Gonzalez, L., Prekeris, R., Steegmaier, M., Advani, R.J., and Scheller, R.H. (1999). SNARE interactions are not selective -Implications for membrane fusion specificity. J Biol Chem 274, 5649–5653.

Yu, H.J., Rathore, S.S., Lopez, J.A., Davis, E.M., James, D.E., Martin, J.L., and Shen, J.S. (2013). Comparative studies of Munc18c and Munc18-1 reveal conserved and divergent mechanisms of Sec1/Munc18 proteins. Proc Natl Acad Sci USA 110, E3271–E3280.

Zhang, X.M., Rebane, A.A., Ma, L., Li, F., Jiao, J., Qu, H., Pincet, F., Rothman, J.E., and Zhang, Y.L. (2016a). Stability, folding dynamics, and long-range conformational transition of the synaptic t-SNARE complex. Proc Natl Acad Sci USA 113, E8031–E8040.

Zhang, Y.L. (2017). Energetics, kinetics, and pathway of SNARE folding and assembly revealed by optical tweezers. Protein Sci 26, 1252–1265.

Zhang, Y.L., and Hughson, F.M. (2021). Chaperoning SNARE folding and assembly. Annu Rev Biochem 90, 581–603.

Zhang, Y.L., Jiao, J., and Rebane, A.A. (2016b). Hidden Markov modeling with detailed balance and its application to single protein folding Biophys J 111, 2110–2124.

Zhao, Y., Fang, Q.H., Herbst, A.D., Berberian, K.N., Almers, W., and Lindau, M. (2013). Rapid structural change in synaptosomal-associated protein 25 (SNAP25) precedes the fusion of single vesicles with the plasma membrane in live chromaffin cells. Proc Natl Acad Sci USA 110, 14249–14254.

Zhou, P., Pang, Z.P.P., Yang, X.F., Zhang, Y.S., Rosenmund, C., Bacaj, T., and Sudhof, T.C. (2013). Syntaxin-1 N-peptide and H_abc_-domain perform distinct essential functions in synaptic vesicle fusion. EMBO J 32, 159–171.

Zorman, S., Rebane, A.A., Ma, L., Yang, G.C., Molski, M.A., Coleman, J., Pincet, F., Rothman, J.E., and Zhang, Y.L. (2014). Common intermediates and kinetics, but different energetics, in the assembly of SNARE proteins. Elife 3, e03348.

