## Supplementary Information for "A dynamic template complex mediates Munc18-chaperoned SNARE assembly"

The SI Appendix contains supplemental text and two supplemental figures.

**Kinetic model of the conformational transitions of the template complex and SNAP-25 binding.** Suppose the template complex undergoes reversible and sequential conformational transitions among the unfolded state, the folded but inactive closed state, and the folded and active open state to which SNAP-25 can bind, as shown in the inset in Fig. 1F with indicated transition rates. Then the probabilities to observe the unfolded template complex $p_u$, the closed template complex $p_c$, the open template complex $p_o$, and the fully assembled SNARE complex $p_s$ at any time $t$ are determined by the Master equations:

$$\begin{cases} \dfrac{dp_u}{dt} = -k_f p_u + k_u p_c \\ \dfrac{dp_c}{dt} = k_f p_u - \left(k_o + k_u\right) p_c + k_c p_o \\ \dfrac{dp_o}{dt} = k_o p_c - \left(k_c + k_b c\right) p_o \\ \dfrac{dp_s}{dt} = k_b c p_o \end{cases} \tag{1}$$

where $c$ is the concentration of SNAP-25 in the solution. Defining the probability vector

$$\boldsymbol{x} \equiv \begin{pmatrix} p_u \\ p_c \\ p_o \end{pmatrix} \tag{2}$$

and the rate matrix

$$\boldsymbol{A} \equiv \begin{bmatrix} -k_f & k_u & 0 \\ k_f & -k_o - k_u & k_c \\ 0 & k_o & -k_c - k_b c \end{bmatrix}, \tag{3}$$

We can rewrite Eq. (1) as

$$\frac{d\boldsymbol{x}}{dt} = \boldsymbol{A}\boldsymbol{x} \tag{4}$$

and

$$p_s\left(t\right) = 1 - \sum_{i=1}^{3} x_i. \tag{5}$$

Suppose the rate matrix has the eigenvalues $\lambda_i, i = 1, \cdots, n$ and the corresponding eigenvectors $\boldsymbol{v}_i$ such that

$$\boldsymbol{AV} = \boldsymbol{VD} \tag{6}$$

where $\boldsymbol{V}$ is a matrix formed by all the eigenvectors

$$\boldsymbol{V} = \left(\boldsymbol{v}_1, \boldsymbol{v}_2, \cdots, \boldsymbol{v}_n\right) \tag{7}$$

and $\boldsymbol{D}$ is a diagonal matrix

$$\boldsymbol{D}=\begin{bmatrix} \lambda_1 & 0 & \cdots & 0 \\ 0 & \lambda_2 & 0 & 0 \\ \vdots & \vdots & \ddots & \vdots \\ 0 & 0 & \cdots & \lambda_n \end{bmatrix} \tag{8}$$

The solution to Eq. (4) is

$$\boldsymbol{x}(t)=\boldsymbol{V}e^{\boldsymbol{D}t}\boldsymbol{V}^{-1}\boldsymbol{x}_0 \tag{9}$$

where $\boldsymbol{x}_0$ is a constant vector whose elements represent the initial state probabilities at time zero.

Given model parameters, we numerically computed the state probabilities according to Eqs. (9) and (5) using MATLAB. These calculations were fit to the experimental measurements to derive the rate constants in the model as best-fit parameters. In our SNAP-25 binding experiments, we held the Syx-VAMP conjugate at a constant mean force close to the equilibrium force of the template complex. In this case, the template complex folded and unfolded with equal rates, which were previously measured, i.e., $k_f = k_u = 3.2$ s$^{-1}$. Correspondingly, the initial state probabilities were chosen as $\boldsymbol{x}_0 = (0.5,\ 0.5,\ 0)$. Thus, our model has three rate constants as fitting parameters. The SNAP-25 binding probabilities were measured at variable SNAP-25 concentration with the fixed time (t=120 s, Fig. 1F) or variable time with fixed SNAP-25 concentration (60 nM, Fig. S1). These probabilities were fit by the model predictions to determine the three rate constants (Fig. 1F). The MATLAB scripts used for these calculations are available upon request.

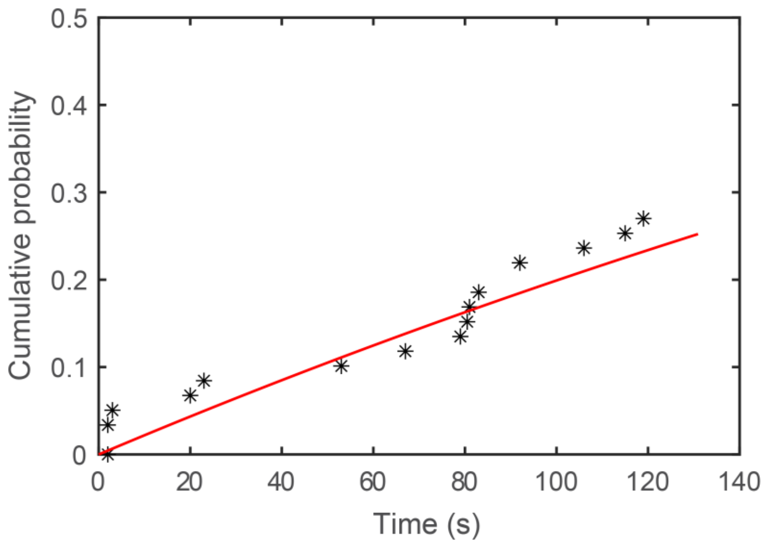


Fig. S1. Cumulative probability calculated from the measured time for SNAP-25 binding in the presence of 60 nM SNAP-25 in the solution (symbols) and its best model fit (red curve).

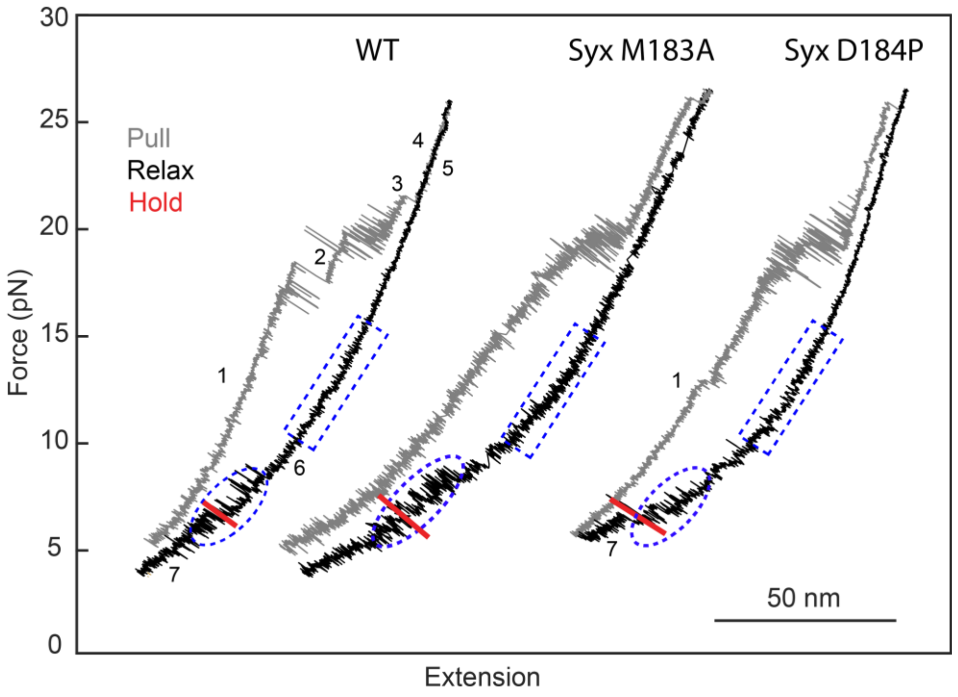


Fig. S2. FECs associated with syntaxin-1 with point mutation M183A or D184P in the linker in the presence of 1 μM Munc18-1 and 60 nM SNAP-25 in the solution. Single pre-assembled SNARE complexes were first pulled to high forces to detect their stepwise unfolding (grey FECs) and then relaxed to low forces to detect reversible refolding of open syntaxin (blue dashed rectangles) and template complexes (blue dashed ovals). Finally, the complexes were held around the equilibrium of the template complex transitions to detect SNAP-25 binding (red regions). The states associated with different FEC regions are labeled as in Fig. 1D.
